# Integrated Plankton and Trace Metal Analysis as Indicators of Organic Pollution Threats from Carp-Cultured Ponds to Adjacent Ecosystems: Implications for Sustainability

**DOI:** 10.1101/2024.01.26.577523

**Authors:** Manasi Mukherjee, Paweł Jarzembowski, Ryszard Polechoński, Jarosław Proćków

## Abstract

Stawy Milickie (Milicz Ponds) is a natural reserve that has a great ecological importance as a Ramsar site and economic importance as one of the largest carp breeding centres in Europe. Continuous anthropogenic pressure on these large ecosystems can lead to a high disturbance on its natural ecology and sustainability. Phytoplankton, one of the most important indicators of any anthropogenic load, especially organic pollution have been studied in the present study. This is the first work ever from Milicz ponds, wherein the diversity, community structure and seasonal variability of phytoplankton with respect to changes in physicochemical parameters and trace metals have been studied. Periodic samples of water and phytoplankton were collected from four selected ponds of various sizes from October 2018 to September 2019. Of the four selected ponds one was unused for carp cultivation. The study reports the presence of 147 phytoplankton species in nine distinct taxonomic groups. The Genera based Algal pollution index indicated presence of organic pollution in both noncultured pond and carp-cultured pond. The heavy metals detected in the concentration order of Cd*>*Mn*>*Fe*>*Zn*>*Cu*>*Pb remained much higher than the safe limits of freshwater aquaculture ponds. The probable sources of these pollutants were identified as artificial feed used in the cultivation of carp and inundation from the river basin. This study emphasizes the urgent need for continuous qualitative and quantitative assessment and monitoring of water parameters, including plankton, in these ponds. This study highlights how the structure of microplankton communities, in conjunction with heavy metal analysis, serves as indicators of the spread of organic pollution in non-cultivated aquaculture environments. It strongly advocates for implementing sustainable measures to safeguard this vital ecosystem.

## 1. Introduction

The ecological health and sustainability of aquatic ecosystems are intricately tied to the diversity and abundance of planktonic organisms. Plankton, which consists of a variety of microscopic organisms, such as phytoplankton, zooplankton, and bacterioplankton, serves as a fundamental component in the balance of aquatic ecosystems. They, especially in aquaculture ponds, play a pivotal role in primary productivity, nutrient cycling, food web dynamics, and oxygen production (Behrenfeld and Boss, 2014). Thus, the community structure characteristics of phytoplankton such as diversity, abundance, and biomass are considered as significant indicators of the quality of an aquatic ecosystem (Kruk et al., 2010). Intricately related to environmental parameters, the monitoring of phytoplankton community structure and diversity is necessary for assessing the health and quality of water (Sabater et al., 2008).

Stawy Milickie (The Milicz Ponds), a complex of interconnected fishponds nestled in the heart of Lower Silesia, Poland, is a crucial aquatic habitat renowned for its ecological significance and biodiversity. These series of ponds that have been used for intensive aquaculture was then established as a nature reserve and protected under the Ramsar Convention in 1963. It covers the ponds of Ruda Milicka, Grabownica, Ruda Sułowska and Radziądz. Covering a large area of 6114.637 ha, it is currently the largest carp breeding centre in Europe (Raftowicz et al., 2021).

A man-made ecosystem turned into natural reserve, Stawy Milickie had encountered escalating anthropogenic pressures and environmental stressors, which posed a potential threat to its fragile ecosystem (Lento et al., 2019). Human-induced factors such as habitat alteration, intensive farming (up to 3.5 t fish/ha), pollution, climate change, and agricultural practices had altered the overall ecological function (Friedel et al., 2008). With the post conservational approach, intense aquaculture has been reduced. However, the lack of any available planktonic studies from this ecosystem makes the understanding of ecological sustainability completely unambiguous. Consequently, understanding the diversity of plankton in Stawy Milickie becomes imperative, not only as a barometer of the health of the ecosystem, but also as a diagnostic tool to gauge the impact of anthropogenic stressors on this vital aquatic habitat.

This study includes the first report of the plankton community, diversity, and abundance of Milicz ponds. In addition to that, the study also reveals the spatiotemporal variations in the structure of the plankton community. Through a comparative analysis between the carp cultured and non-carp cultured Milicz ponds, the study also establishes the sustainable conditions of the Milicz ponds. The results of the study indicate that the ponds used for carp culture have a healthy and sustainable environment due to regular drying and catchment from the Barycz River. The study thus aims to offer a foundation for informed conservation strategies, helping to preserve and restore this unique aquatic ecosystem.

## 2. Methods

### 2.1. Study area

The study focused on four distinct ponds within the Ruda Sułowska complex utilised for aquaculture purposes, receiving periodic supplementation with artificial fish feed. To delineate differences in plankton diversity between carp cultured and non-carp cultured, Pond 1, a narrow channel void of cultivation due to its small size, was selected. Ponds 2, 3, and 4, utilised for fish cultivation, were chosen based on progressively larger sizes. However, Ponds 1, 2, and 3 were among those completely dried for pest control measures during March 2019, allowing plankton sampling solely from Pond 4 in April. The essential details on the coordinates and area of each selected pond are presented in Table 1.

**Table 1:**
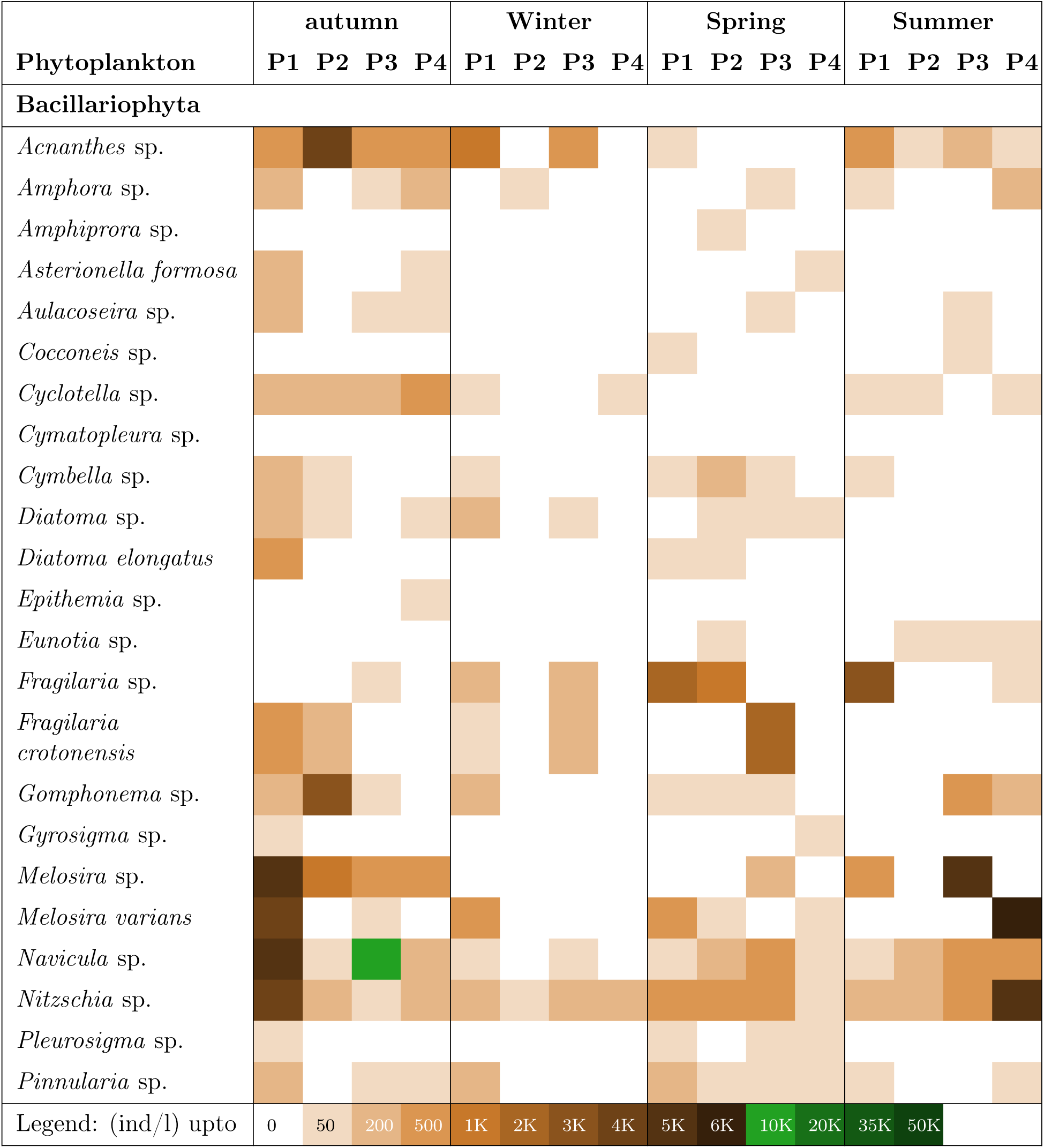
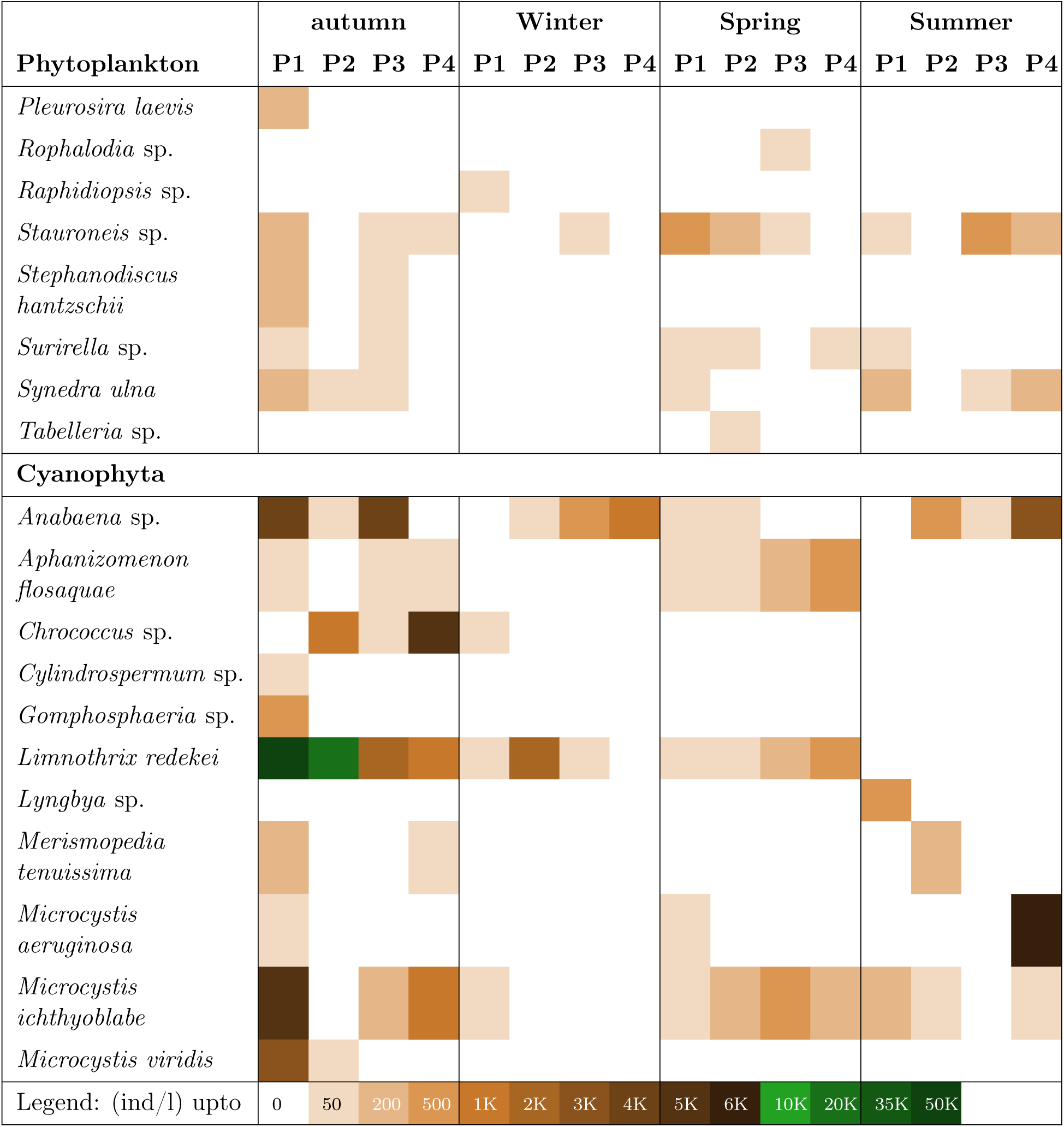
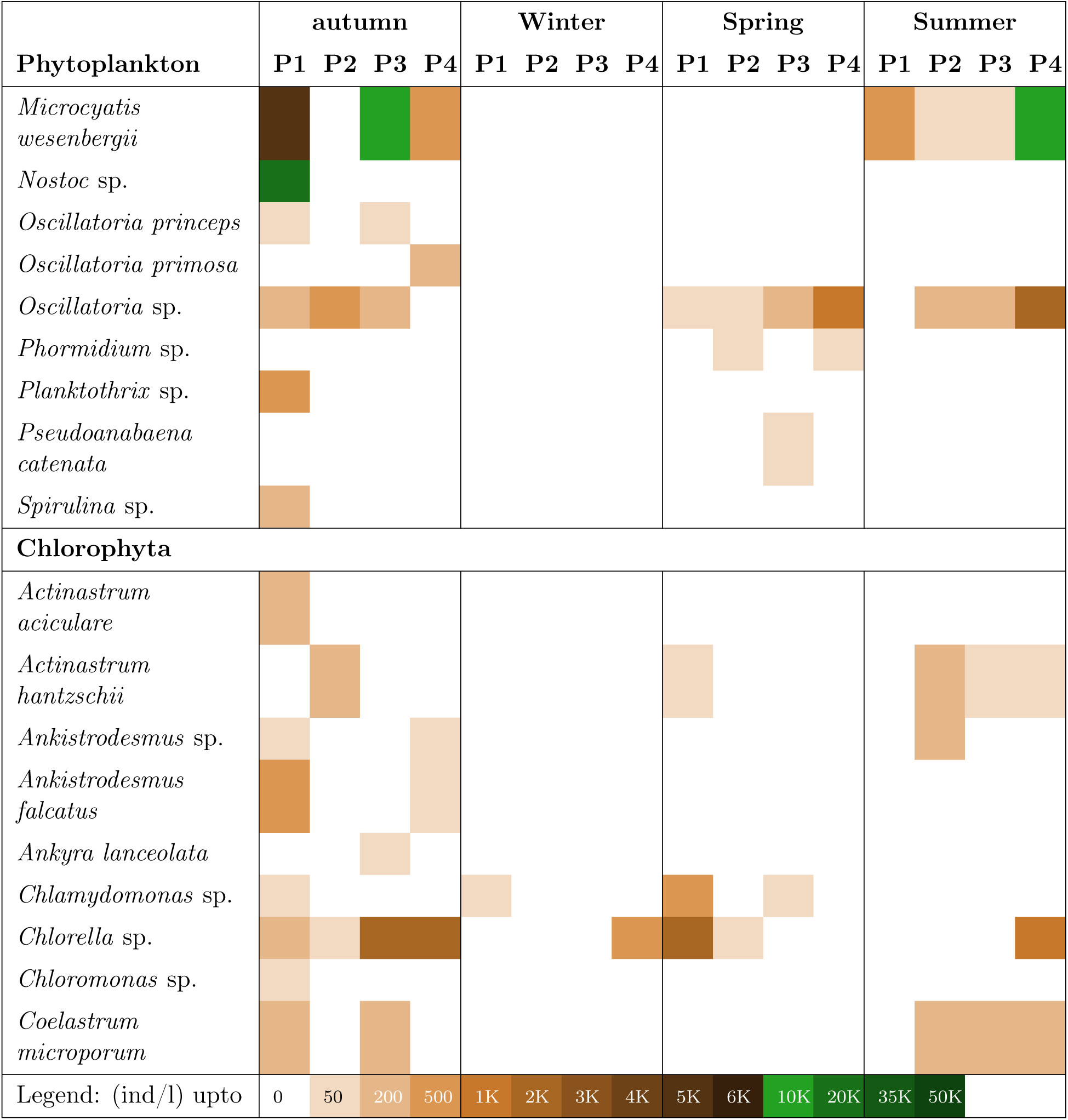
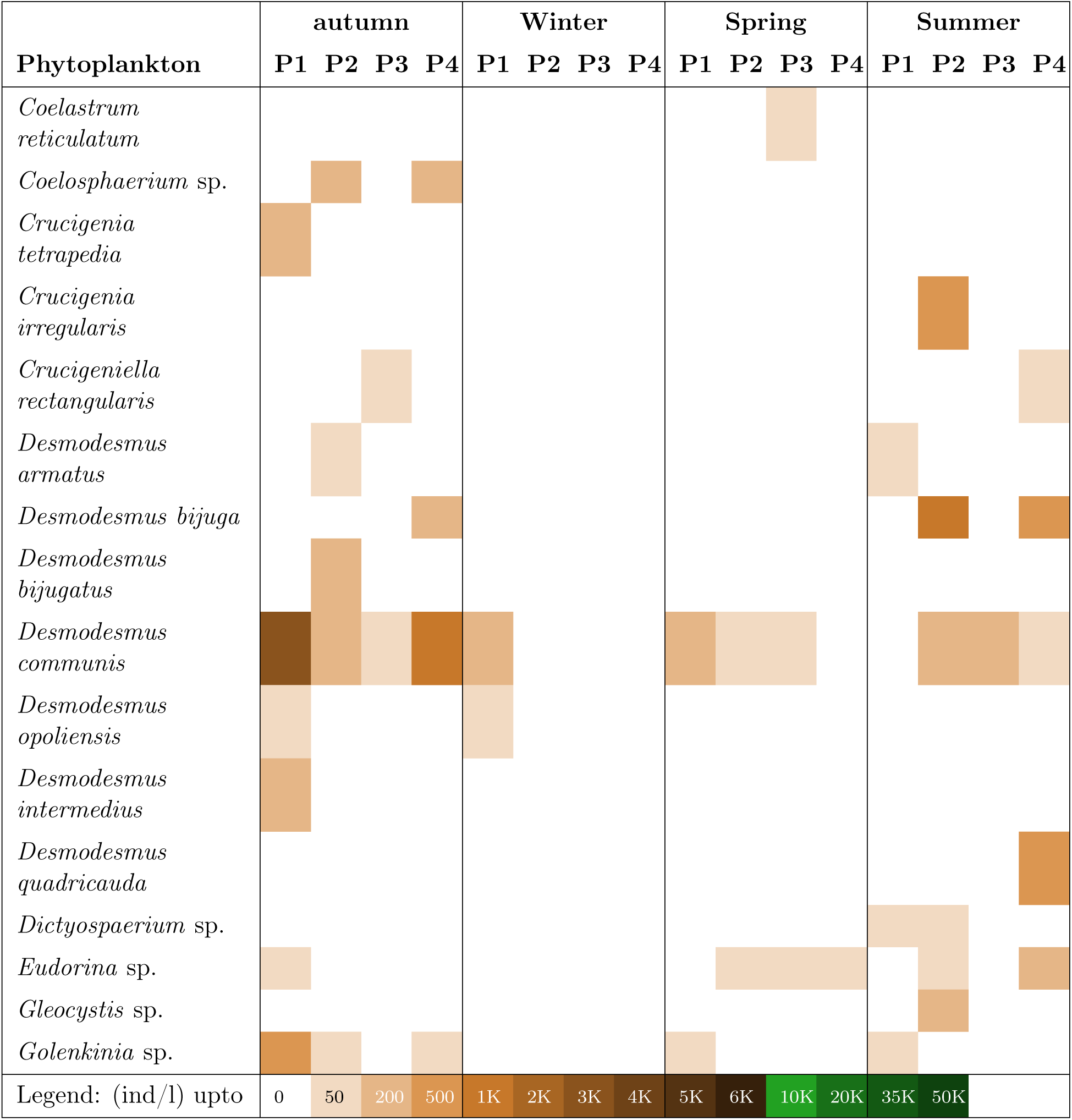
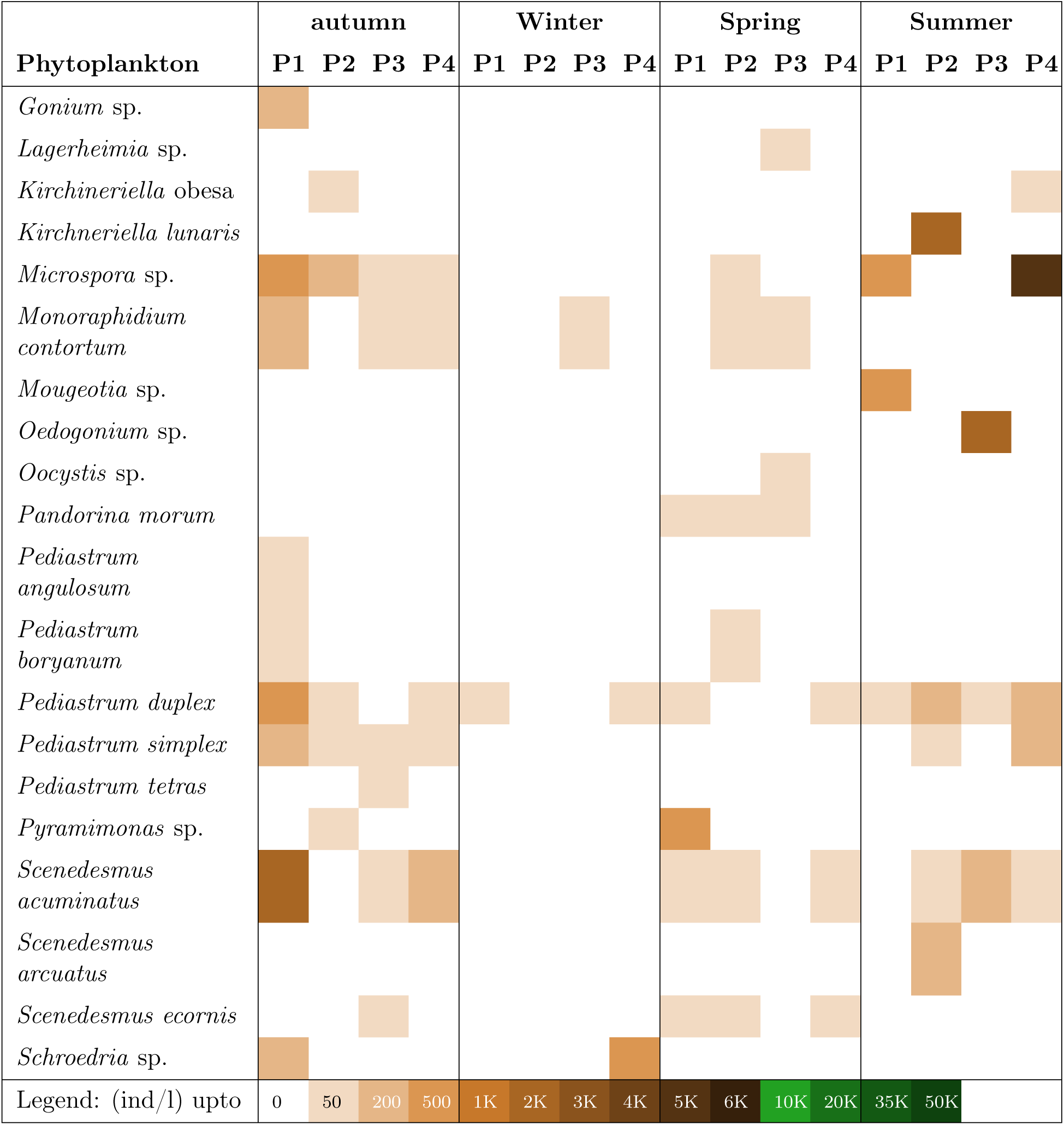
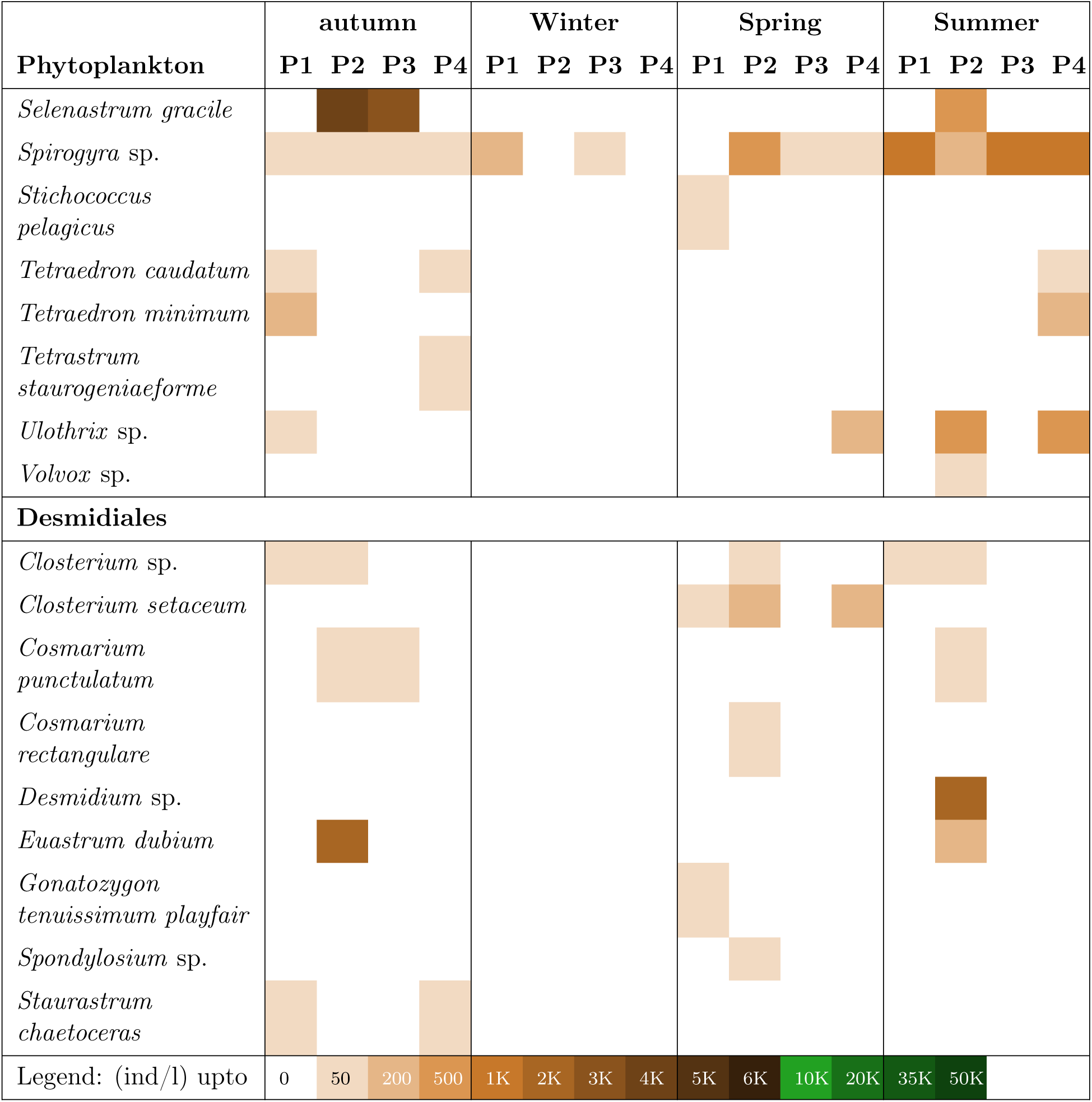
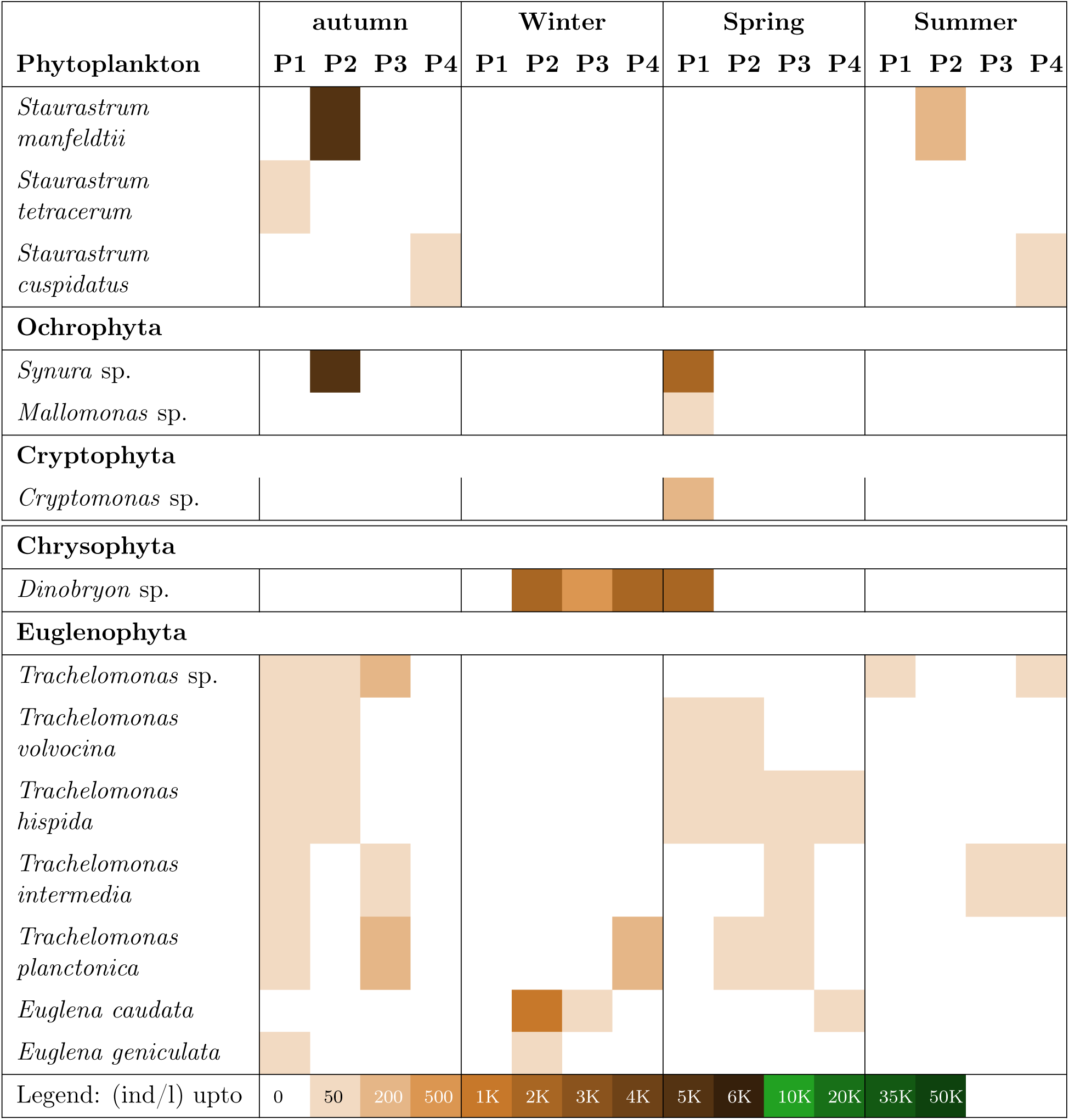
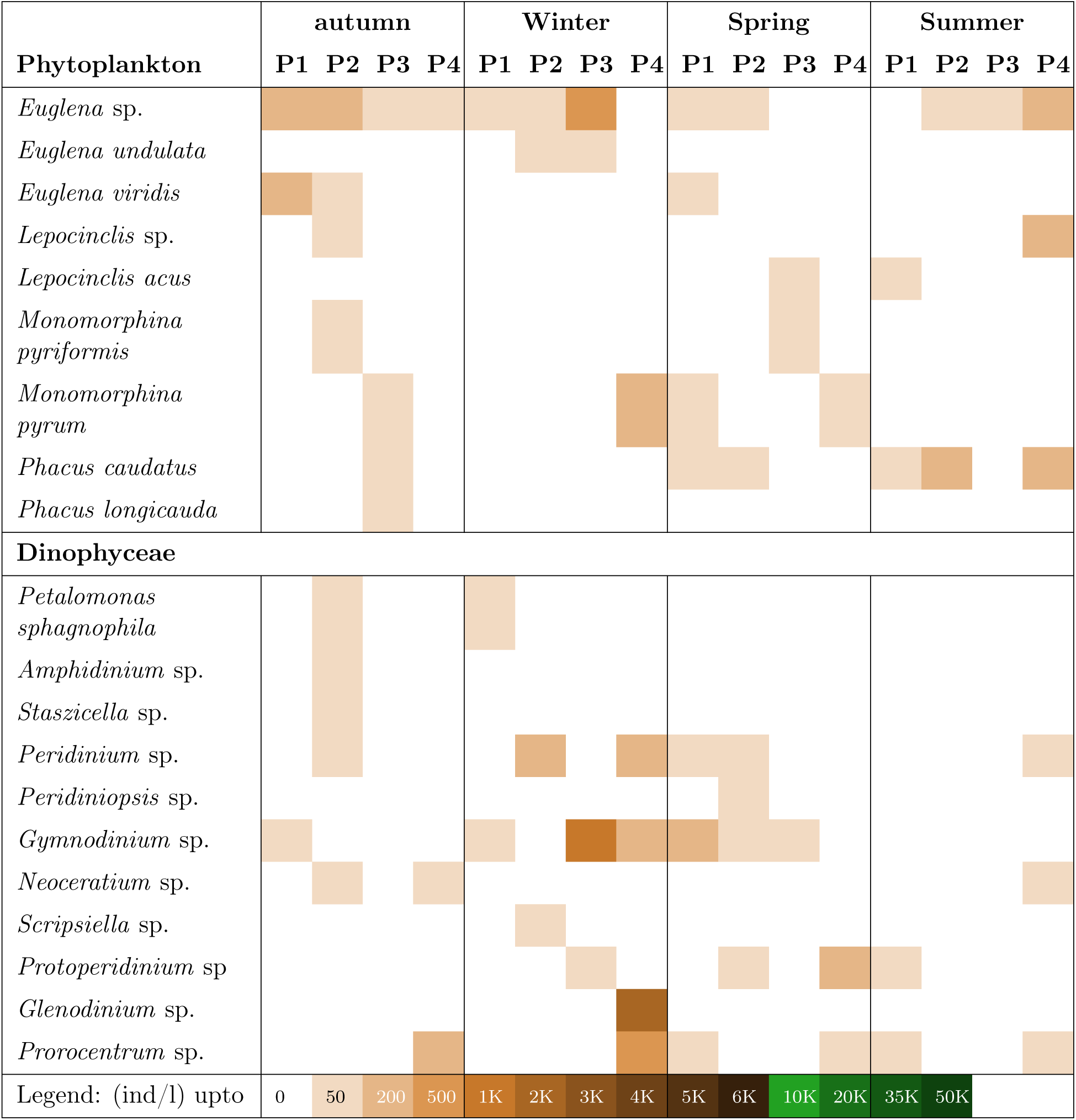
Seasonal variation in the diversity and abundance of phytoplankton in Milicz ponds.

**Table 2:**
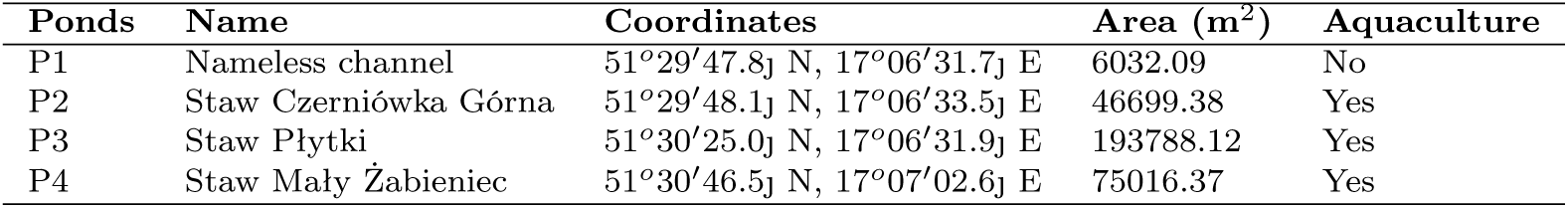
Details of sampled ponds of Stawy Milickie (Milicz ponds)

| **Ponds** | **Name** | **Coordinates** | **Area (m<sup>2</sup>)** | **Aquaculture** |
| --- | --- | --- | --- | --- |
| P1 | Nameless channel | 51°29′47.8j N, 17°06′31.7j E | 6032.09 | No |
| P2 | Staw Czerniówka Górna | 51°29′48.1j N, 17°06′33.5j E | 46699.38 | Yes |
| P3 | Staw Płytki | 51°30′25.0j N, 17°06′31.9j E | 193788.12 | Yes |
| P4 | Staw Mały Żabieniec | 51°30′46.5j N, 17°07′02.6j E | 75016.37 | Yes |

### 2.2. Sample collection and data analysis

#### 2.2.1. Plankton

Plankton sampling was carried out from October 2018 to April 2019 using a 25 micrometer plankton net, filtering 100 litres of water from each sampling site. Collection during November and January involved breaking the ice layer to retrieve samples due to complete freezing of the ponds. The filtered samples were appropriately labelled, preserved using Lugol’s solution, and stored in a laboratory refrigerator for subsequent analysis. Sample examination and enumeration were performed using an Opta-Tech MB200 compound microscope equipped with a Delta Optical DLT-Cam PRO, an 18MP camera. Taxonomic identification up to the genus or species level relied on established keys (Taylor, 1976; Tomas and Hasle, 1997; Cremer et al., 2007; Faust and Gulledge, 2002)

Although all four ponds are subjected to periodic drying for pest control and subsequent feeding from the Barycz River, Pond 1 differs in that it is not used for carp culture and does not receive artificial feed. Consequently, Pond 1 served as the control in this investigation, sharing similar environmental conditions with the other ponds, except for the absence of carp cultivation. As Pond 1 remained significantly smaller, varying pond sizes were strategically selected to account for physical constraints related to the area in this study.

Diversity indices like Shannon, Dominance, Margalef Richness and Evenness were estimated using Past software.

For identifying organically polluted ponds, Palmer (1969) Algal Generic Pollution Index was employed.

#### 2.2.2. Environmnetal parameters

Physicochemical analysis of water was carried out using standard methods (APHA, 2005). Cd, Pb, Ni, Cu, and Zn concentrations were determined using an atomic absorption spectrophotometer (SpectrAA FS220, Varian, Australia).

## 3. Results and Discussion

Investigating the diversity of plankton within Stawy-Milickie has revealed a wide range of phytoplankton species, totalling 147 in nine distinct taxonomic groups. These encompass 33 bacillariophytes, 18 cyanophytes, 53 chlorophytes, 17 euglenophytes, 12 desmidiales, 10 dinophytes, 2 Ochrophyta, 1 Chrysophyceae, and 1 Cryptophyta. A detailed distribution of these identified phytoplankton species in the four studied ponds during different seasons is outlined in Table 1.

Throughout the duration of the study, Pond 1 consistently exhibited the highest phytoplankton diversity and abundance (Fig.2). The phytoplankton Species richness (S) and abundance displayed a pattern of increase in autumn and winter while decreasing in winter and summer. Pond 1 of all the ponds displayed the highest species richness and abundance of phytoplankton during all seasons except summer.

**Figure 1:**
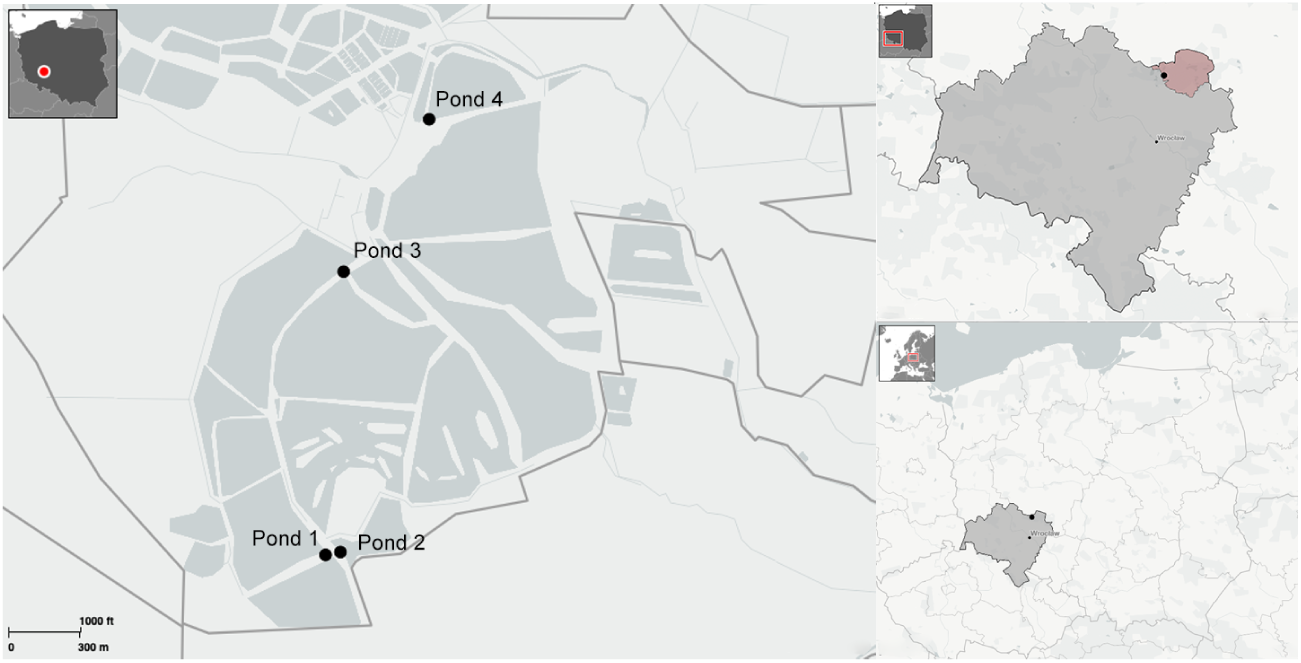
Illustration depicting the spatial distribution of four designated Milicz ponds denoted by black dots and labeled as Pond 1, 2, 3, and 4. These ponds located in Poland (depicted in dark grey on the left top side) serve as representatives of the Milicz reserve, situated within the Milicz powiat (highlighted in red), which is part of the Lower Silesian voivodeship (marked in grey).

**Figure 2:**
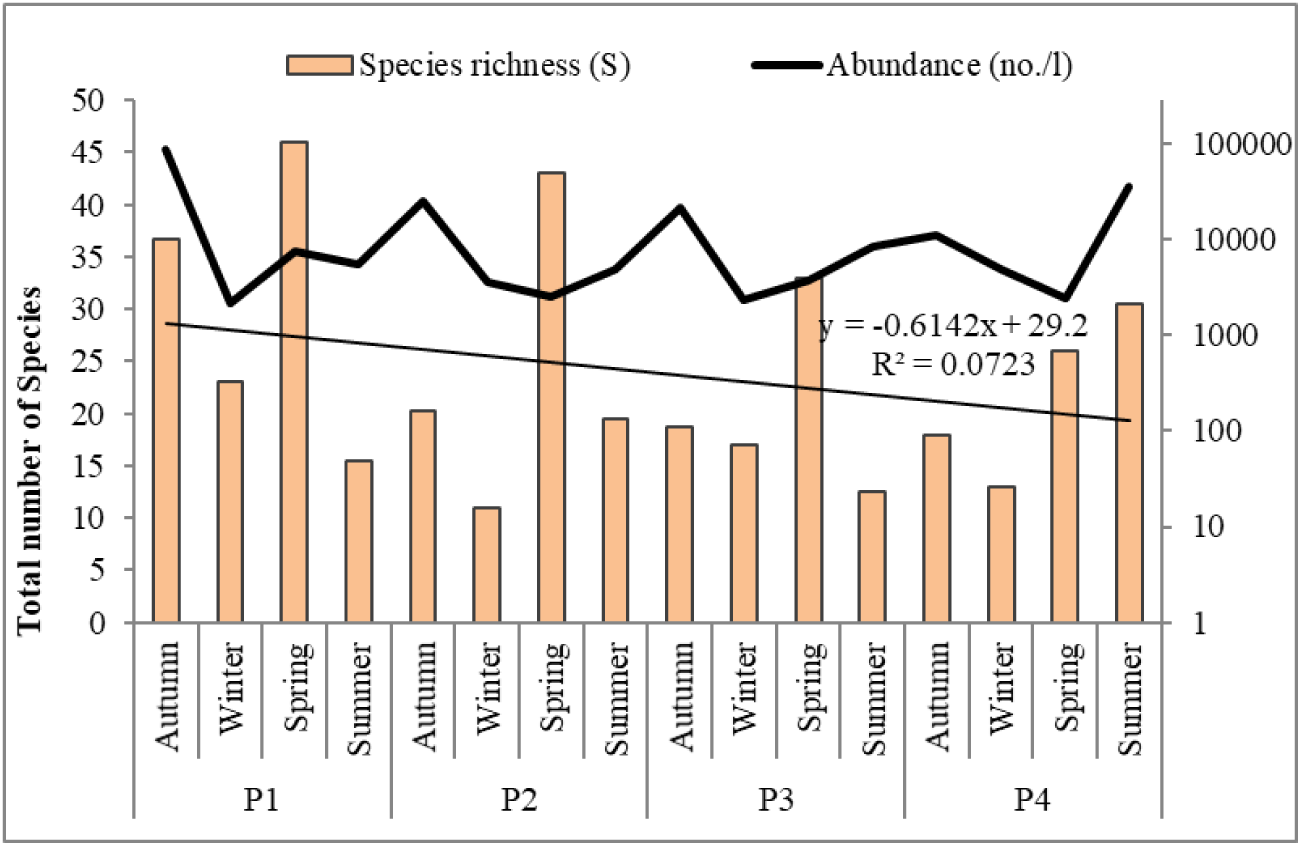
Representation of seasonal and site wise changes in phytoplankton Species richness (S) and abundance. The linear trendline shows a gradual decrease in phytoplankton richness and abundance in Pond 1 to 4 during autumn and summer.

Seasonal fluctuations were evident, with certain species prevalent in specific seasons, yet absent in others. For example, species such as *Aphanizomenon flos-aquae*, *Microcystis* spp., *Oscillatoria* spp., *Desmodesmus* spp., *Scenedesmus* spp., *Monoraphidium contortum*, and *Trachelomonas* spp. were dominant in various seasons but absent during winter. On the contrary, *Dinobryon* sp., a chrysophyte, only occurred during winter. *Limnothrix redekei* exhibited significantly higher abundance during autumn in all four studied ponds, notably contributing (132563 individuals per litre) to the highest abundance of phytoplankton in Pond 1 during that month. However, its abundance gradually decreased in both winter and spring.

As the difference between Milicz carp cultured and non-carp cultured ponds was evident, the ponds were analysed for their physicochemical properties to identify any specific variation (Table 3). The water temperature of all the ponds remained relatively similar ranging between 9 to 11*°*C in autumn, while 1 to 2 *°*C during winter. Parameters such as pH and carbonate hardness of all the ponds during both seasons remained within the favourable limit of aquaculture ponds (Wurts and Durborow, 1992). However, it is important to note that PO4 (%) remained considerably high and NH3 very low in Pond1 during both seasons. As mentioned by Durborow et al. (1997), phosphorus promotes algae growth that utilizes nitrogen, thus reducing NH3. Therefore, this explains the comparatively higher abundance of phytoplankton in Pond 1 throughout the year.

**Table 3:** Physicochemical properties of Milicz ponds during the study period.

| Seasons | Ponds | Temp.<br>(°C) | pH | Conduc.<br>(uS/cm) | KH<br>(mg/l) | NH3<br>(mg/l) | NO2<br>(mg/l) | NO3<br>(mg/l) | PO4<br>(mg/l) | SO4<br>(mg/l) |
| --- | --- | --- | --- | --- | --- | --- | --- | --- | --- | --- |
| **Autumn** | **P1** | 11 | 7.25 | 396 | 142.8 | 0.02 | 0.01 | 5 | 0.2 | 10 |
|  | **P2** | 9 | 7.42 | 352 | 107.1 | 0.1 | 0 | 0 | 0.05 | 10 |
|  | **P3** | 9.5 | 7.62 | 341 | 124.9 | 0 | 0.015 | 5 | 0 | 10 |
|  | **P4** | 10 | 7.64 | 373 | 142.8 | 0.2 | 0.002 | 3 | 0.15 | 15 |
| **Winter** | **P1** | 1 | 7.15 | 457 | 160.6 | 0.01 | 0.02 | 10 | 0.25 | 10 |
|  | **P2** | 1 | 6.94 | 452 | 142.8 | 0.25 | 0 | 5 | 0.05 | 15 |
|  | **P3** | 2 | 6.97 | 395 | 160.6 | 0 | 0.02 | 2 | 0 | 10 |
|  | **P4** | 1 | 7.12 | 380 | 142.8 | 0.5 | 0.025 | 5 | 0.05 | 25 |

Figure 3 illustrates the proportional representation of various phytoplankton groups studied in all ponds. The Spatio-temporal dynamics revealed a shift in the composition of the phytoplankton community across all ponds with changing seasons. The figure also corroborates the results from Fig. 2 that the overall phytoplankton abundance remained higher during the autumn seasons, whereas it dropped during spring, summer and winter.

**Figure 3:**
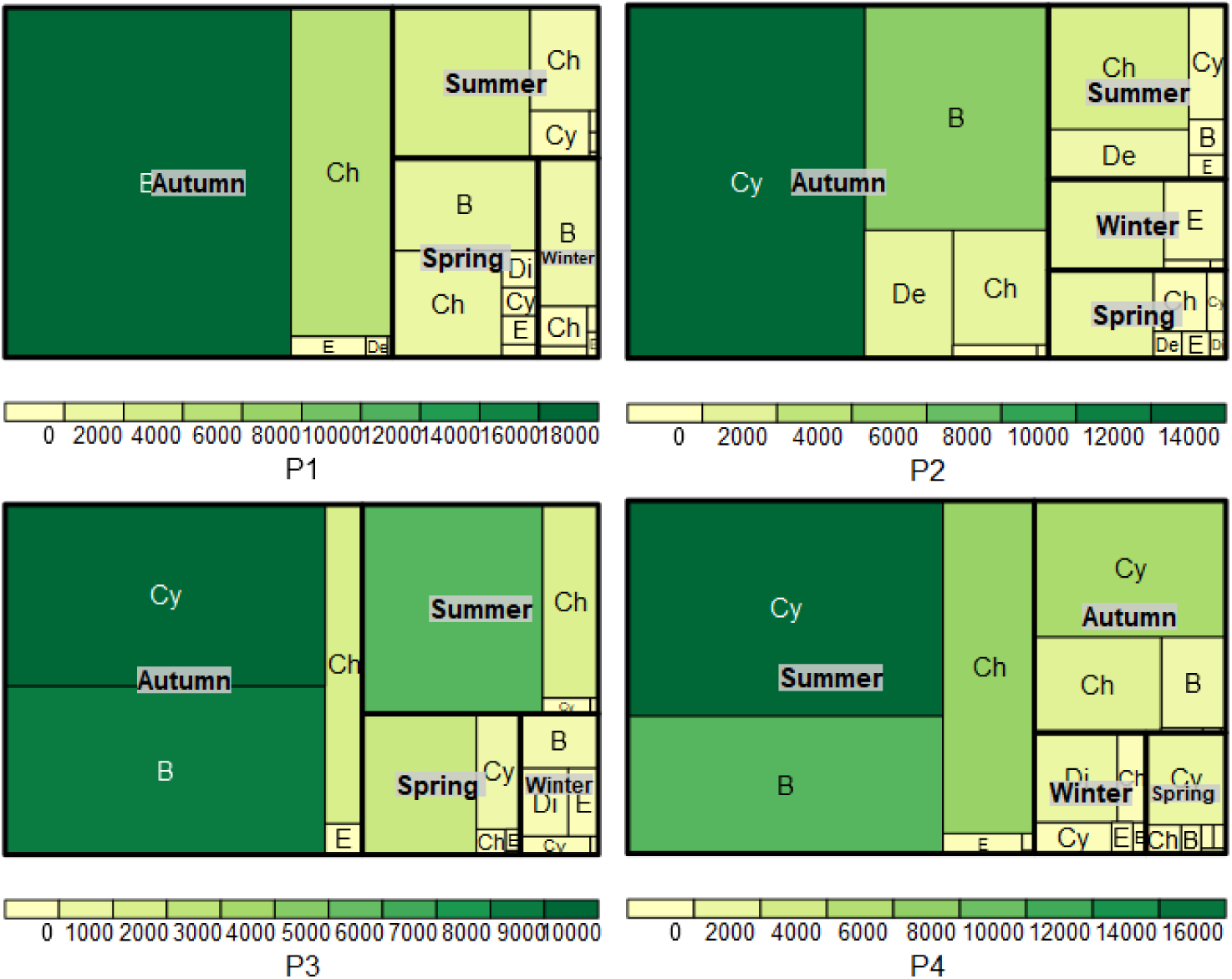
Mean seasonal abundance of the identified phytoplankton groups from all the four sampling sites (Ponds: P1, P2, P3 and P4) of Stawy Milickie. The thickest black outline of each rectangle of the treemap corresponds to each season (autumn, winter, spring and summer). Each rectangle within the seasonal rectangle corresponds to the proportion of a specific phytoplankton group (B: Bacillariophyta. Cy: Cyanophyta, Ch: Chlorophyta, De: Desmidiales, E: Euglenophyta and Di: Dinophyta). The gradient of the colour of each rectangle is based on the corresponding abundance value of the phytoplankton group.

Among these groups, Bacillariophytes and Cyanophytes consistently emerged as the primary dominants in all ponds. However, Pond 1 was the only site that exhibited a greater prevalence of Bacillariophytes throughout all the seasons. Bacillariophytes have also been studied to have the potential to store nitrate intracellularly for Dissimilatory Nitrate Reduction to Ammonium (DNRA) (Stief et al., 2022). This therefore explains the higher abundance of Bacillariophytes and the comparatively higher concentration of nitrate, nitrite and phosphate in Pond1. A higher abundance of Bacillariophytes as is found in Pond 1 is an indicator of organic pollution (Arumugham et al., 2023; Khalil et al., 2021). On the other hand, the variation in seasonal community composition of phytoplankton from Ponds 2, 3 and 4 showed higher occurrences of Cyanophytes and Chlorophytes. High nutrient loads from aquaculture can promote the production of organic matter. Then the decomposition and release of regenerated ammonium (in P2, P3 and P4) provide a favourable environment for the growth of Cyanobacteria in autumn and summer (Newell et al., 2019). As is known from the ice-covered natural fresh water ecosystems (Lento et al., 2019), the seasonal prevalence of Euglenophytes and Dinophytes was higher during winter from all the ponds, Dinophytes especially spiking in Pond 3 and 4. Desmidiales remained higher in Pond 2. A higher abundance of Chrysophytic and Ochrophytic flagellates such as Dinobryon sp. and Synura sp. during winter and winter suggested an oligo-mesotrophic condition prevailing in the studied ponds.

This was further validated with diversity indices like Shannon (Fig. 4a), Dominance (Fig. 4b), richness (Fig. 4c) and Evenness (Fig. 4d). A Shannon-Wiener index (H‘) less than 1.0 is considered polluted for aquaculture ponds (Pleto and Cabillon, 2022). Except Pond 3, the Shannon index ranged above 1, indicating that there was no significant pollution. Pond 1 showed a comparatively lower value of Simpson’s dominance (Fig. 4b) and a higher value of Margalef’s species richness (Fig. 4c), signifying reduced species dominance and greater evenness (Fig. 4d) for a more diverse phytoplankton population to flourish. Research indicates that regardless of fluctuations in absolute environmental temperatures, phytoplankton growth rates tend to increase with greater species richness.

**Figure 4:**
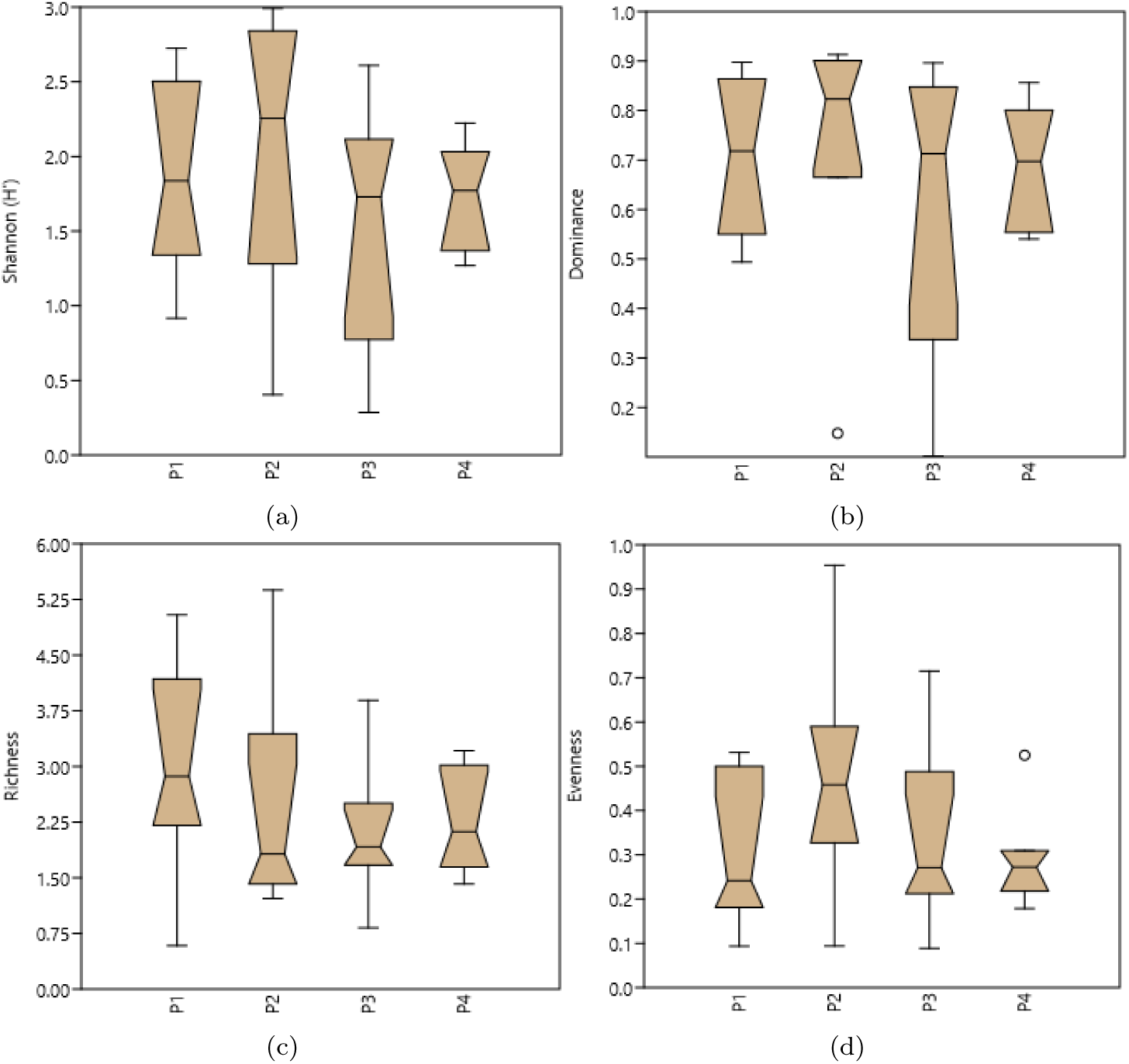
Biodiversity indices of four studied ponds. Shannon index (a), Dominance (d), Margaleffs Richness (c) and Evenness (d) from ponds P1, P2, P3 and P4. The upper and lower quartiles are measured, excluding the outliers. Pond 4 of all shows the least variability in all the indices.

According to the diversity indices, the ponds do not show significant organic pollution, whereas the composition of the plankton community indicated the presence of organic pollution, especially in Pond 1. Thus, the Generic Algal Pollution Index which is based on the top 20 pollution indicating genera and their respective scores was estimated to understand the extent of pollution in the ponds, if any. Figure 5 indicates that Ponds 1 and 4 have high organic pollution, whereas Ponds 2 and 3 have moderate organic pollution. The status of the pollution varies seasonally, however, indicating more organic pollution during autumn and summer and less organic pollution during winter and spring.

**Figure 5:**
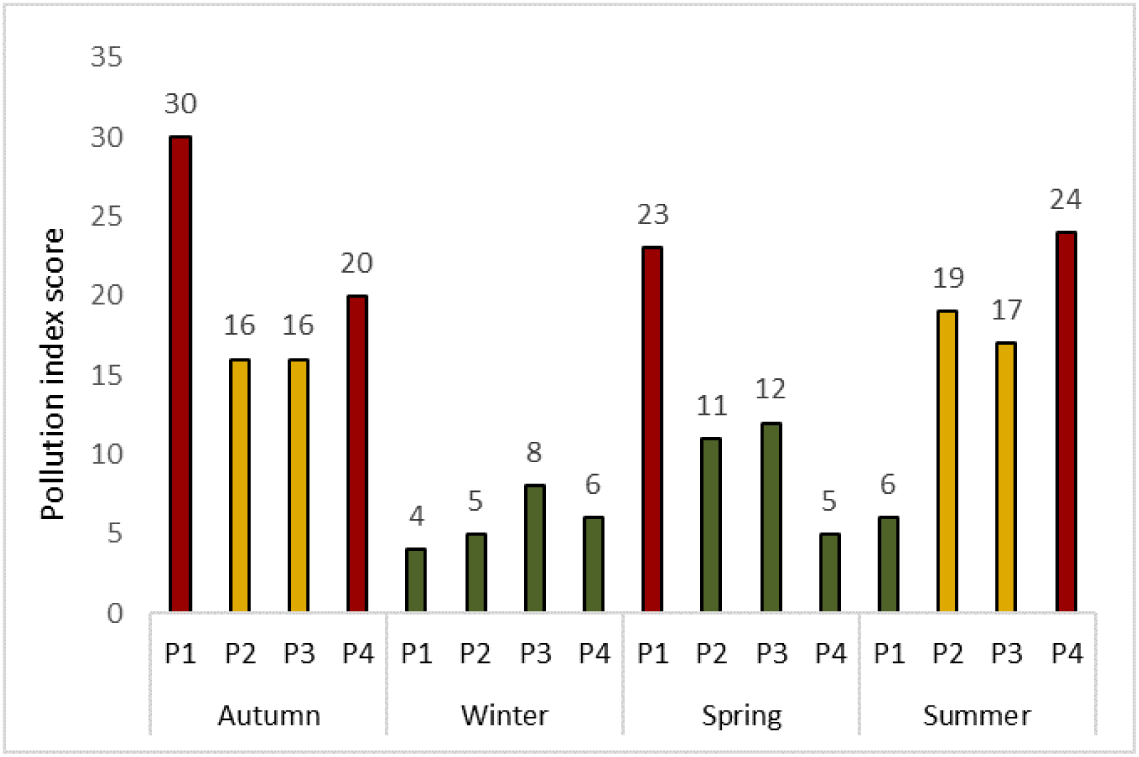
Genera based Algal pollution index as described by Palmer (1969) to estimate organic pollution of Milicz ponds. Score *>*20 indicate high organic pollution; represented in red. Scores = 15-19 indicate moderate organic pollution, represented with yellow and Score: *<*15 indicate low organic pollution, represented with green. The seasonal assessment indicates that the organic pollution is lowest in winter and spring, whereas it is highest during autumn and summer. Ponds 1 and 4 show high organic pollution, whereas Ponds 2 and 3 show moderate organic pollution.

The ponds were thus analysed for their trace metals to understand the basis of organic pollution and their causing factors. The results showed the presence of trace metals Cd, Mn, Fe, Zn, Cu, Pb in the concentration order of Cd*>*Mn*>*Fe*>*Zn*>*Cu*>*Pb (Table4). Pokorny et al. (2015) showed the presence of these trace metals from other Milicz ponds except Mn and Fe. All trace elements recorded were much higher than the safe limits of heavy metal concentrations for an aquaculture pond, especially during autumn and summer.

A discussion with the farming community revealed that the carp cultured ponds are artificially fed with feed like wheat, triticale, corn and rye. The feeding occurs from June to September and occasionally in October. The feeding coefficient depends on the type of feed. The food factors for the feed type are: wheat = 5, rye = 7, and barley = 5. Food factor = amount of food consumed per 1 kilogram of increment. Artificial feeding is known to cause heavy metal pollution in cultured ponds (Rahman and Ghosh, 2021; Hossain et al., 2023). Thus, in the present study it induces the possibility that the treatment of cultured ponds with artificial feed during autumn and summer has led to the development of organic pollution in all ponds. It is also important to note that Pond 1 though is a noncultured pond and is not treated with artificial feed, it shows high trace metal value (Table 4) as well as pollution index (Fig. 5). There could be two possibilities for this outcome. Primarily the inundation of ponds from catchment (Barycz river) can lead to the influx of heavy metals overflowing from other ponds to such noncultured ponds. The second possibility is that the Barycz river itself could be a point source of heavy metal pollution (Sojka and Jaskuła, 2022). Moreover, as these noncultured ponds do not undergo the periodic drying and refilling process, the accumulation of pollutants remains higher causing severe effects on its biota and sustainability. However, while the other ponds were found to have similar amounts of trace metals, the organic pollution remained lower due to a periodic drying and cleaning processes. However, the trace element concentrations exceeding safe limits and moderate organic pollution expressed by the pollution index point towards a probably unsafe environment for biota in these cultured ponds.

**Table 4:**
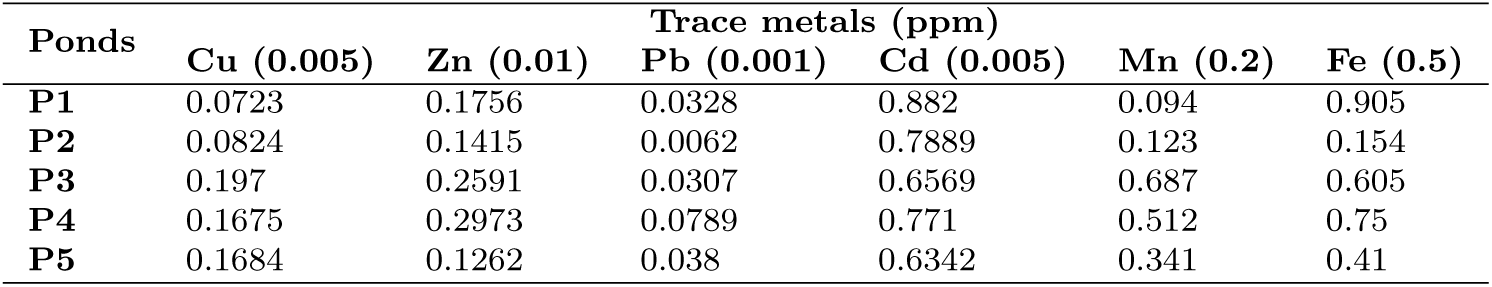
Metal concentrations (ppm) in selected Milicz pond water. The maximum safe concentration for freshwater is given in parentheses based on Boyd (2009)

| **Ponds** | **Trace metals (ppm)** |  |  |  |  |  |
| --- | --- | --- | --- | --- | --- | --- |
|  | **Cu (0.005)** | **Zn (0.01)** | **Pb (0.001)** | **Cd (0.005)** | **Mn (0.2)** | **Fe (0.5)** |
| **P1** | 0.0723 | 0.1756 | 0.0328 | 0.882 | 0.094 | 0.905 |
| **P2** | 0.0824 | 0.1415 | 0.0062 | 0.7889 | 0.123 | 0.154 |
| **P3** | 0.197 | 0.2591 | 0.0307 | 0.6569 | 0.687 | 0.605 |
| **P4** | 0.1675 | 0.2973 | 0.0789 | 0.771 | 0.512 | 0.75 |
| **P5** | 0.1684 | 0.1262 | 0.038 | 0.6342 | 0.341 | 0.41 |

## 4. Conclusion

The phytoplankton diversity, community composition and algal pollution index of Milicz ponds under study show the presence of moderate to high organic pollution. This outcome is supported by the level of physicochemical parameters like NH3, NO3 and NO2. The trace elements detected in the pond waters also corroborate the results, indicating the possibility of the spread of trace metal pollution through the feed or catchment as the point source. Therefore, the outcome of the present study suggests monitoring both artificial feed and the quality of water in the Milicz pond catchment to determine measures for the sustainability of this ecosystem.

## Acknowledgements

We express our gratitude to the farming community of Milicz Ponds for their valuable contributions during discussions.

## Authorship contribution statement

**Manasi Mukherjee** contributed in study conception and design, data collection, analysis, interpretation of results and writing. **Paweł Jarzembowski** and **Ryszard Polechoński** contributed to data collection, analysis and interpretation of physicochemical data. **Jarosław Proćków** contributed to supervision, writing, reviewing and editing.

